# A lipid acyl code-based Dip2-Pkc1 signalling axis maintains mitochondrial integrity in eukaryotes

**DOI:** 10.64898/2026.07.21.739733

**Authors:** Santhosh Kumar, Sakshi Shambhavi, Azeen Zehra, Arnab Chakraborty, Suhail Madhar Hanif Saleem, Abhijit Mohapatra, Biswajit Pal, Shashi Vardhan Kalivendi, Siddhesh S. Kamat, Rajan Sankaranarayanan

## Abstract

Organelle membranes employ diverse lipids to relay key signals for efficient coordination of cellular processes. Diacylglycerol (DAG) is a simple yet critical lipid secondary messenger, but the regulatory mechanisms and functional implications for its distribution remain poorly understood. We have recently shown that Protein Kinase C (Pkc1) activation is driven by selective DAGs (C36:0, C36:1), whose levels are governed by Disco-interacting protein 2 (Dip2) (Shambhavi et al., 2025). Here, through genetic, chemical, and lipidomic screens, we show that the absence of Dip2 leads to specific DAG accumulation on the mitochondrial membrane and impacts its morphology, function, and quality control in yeast. Remarkably, the elevated DAGs in *Δdip2* promote translocation of Pkc1 to mitochondria via its DAG-binding C1 domain, but not the HR1 domain, suggesting functional partitioning between the regulatory domains. We also show that only specific DAGs, not the bulk DAGs, are required for Pkc1’s targeting and inactivating Phospholipase C (Plc1) restores Pkc1 localisation and the associated mitochondrial defects. In addition, we identify that respiratory growth triggers specific DAG (C36:1) accumulation in the mitochondria, thereby promoting Pkc1 recruitment. Furthermore, we establish that the Psi1-Plc1-Dip2 axis is required for survival under respiratory growth conditions. Taken together, our study uncovers a novel, Dip2-mediated, unconventional Pkc1 signalling axis for maintaining mitochondrial homeostasis under nutrient transition and highlights how distinct lipid fingerprints enable precise control over organellar homeostasis.

## Introduction

Spatial segregation of biochemical reactions into membrane-bound organelles is a defining feature of eukaryotic cells and lipids constitute a major portion of these membranes (Behnia & Munro., 2006; van Meer et al., 2008). Beyond their crucial role in membrane biogenesis and maintaining its integrity, several bioactive lipids act as secondary messengers in signal transduction either by recruiting or activating effector proteins. Numerous studies have elucidated the role of different lipid classes in critical signalling processes – for example, Phosphoinositides in cell polarity, Diacylglycerol and Phosphatidic acid in mitochondrial dynamics, Sphingosine 1-phosphate in cell differentiation etc., (Ryu et al., 2006; Adachi et al., 2016; Xie & Bankaitis., 2022; Pemberton et al., 2025) and these functions have largely been attributed to the differences in their head group. However, the regulatory potential of fatty acyl tails in these processes is an emerging yet underexplored theme. Recently, a few studies have highlighted that variations in the chemical nature of fatty acyl tails, i.e., chain length and degree of unsaturation, can significantly modulate the signalling pathways (Schuhmacher et al., 2020; Mao et al., 2023; Qiu et al., 2024; Kchir et al., 2026).

We have earlier uncovered the role of a conserved eukaryotic protein family, DISCO-Interacting Protein 2 (Dip2), as a homeostatic regulator of specific subsets of diacylglycerols (DAGs) by converting them to triacylglycerols (TAGs) (Mondal et al., 2022). We have shown that Dip2 knockout leads to constitutive activation of unfolded protein response (UPR) pathway and sensitivity to tunicamycin-induced ER stress. We have demonstrated that Dip2 is required for proper vacuolar membrane fusion and osmoadaptation, as altered levels of Dip2 results in aberrant vacuolar morphology. Interestingly, we also found that Dip2 localises primarily to mitochondria and associates with the vacuole via mitochondria-vacuole contact sites. Thus, the study identified a previously unknown route of DAG metabolism and established Dip2 as a new family of acyl chain-specific DAG regulator.

In our recent study, we have identified Dip2 as a regulator of Protein Kinase C (Pkc1), one of the well-known lipid-based signalling modules in eukaryotes (Shambhavi et al., 2025). We showed that Dip2-regulated specific DAGs (C36:0, C36:1) are required for Pkc1 activation in *Saccharomyces cerevisiae*. Through in vitro and in vivo experiments, we showed that Pkc1 is activated only by the specific DAG pool regulated by Dip2 and not by the bulk DAGs. These Dip2-regulated DAGs are sourced from acyl chain remodelling of phosphatidylinositols (PI) by Psi1, followed by Phospholipase C (Plc1)-mediated hydrolysis of phosphoinositides. Furthermore, through extensive co-evolutionary analysis, we established that Dip2 and Pkc1 have co-emerged and co-evolved, marking the emergence of a primitive DAG-based signalling axis in eukaryotes.

Pkc1, a serine-threonine kinase, is an essential signalling protein that regulates various aspects of cellular physiology, including differentiation, cell migration, proliferation, angiogenesis, neuronal branching, cell wall integrity etc. (Newton., 2018). Consequently, dysregulated Pkc1 signalling is implicated in several pathologies like diabetes, Alzheimer’s, heart disease and various carcinomas (Etcheberrigaray et al., 2004; Palaniyandi et al., 2009; Wallace et al., 2014; Callender & Newton., 2017; Isakov., 2018; Marrocco et al., 2019; Martin-Perez et al., 2020). Therefore, understanding the intricate mechanisms of Pkc1 regulation is of utmost importance. Given the establishment of Dip2-Pkc1 signalling axis at the base of eukaryotic evolution, key questions remain regarding the mechanistic basis of their crosstalk. What is more intriguing to resolve is how Dip2, a protein localised at the mitochondrial membrane, regulates Pkc1, which localises at the bud site or bud neck in yeast (Denis & Cyert., 2005). Moreover, the subcellular localisation of the selective DAGs regulated by Dip2 and how their spatial distribution shapes Pkc1 activity and its downstream processes remain unexplored.

Here, we show that Dip2-mediated DAG regulation occurs at the mitochondrial membrane and is critical for maintaining proper mitochondrial morphology and function in yeast. Absence of Dip2 (*Δdip2*) leads to the translocation of Pkc1 to mitochondria, a process dependent on its DAG-binding C1 domain. Through genetic and chemical screens, we show that only Dip2-regulated specific DAGs, not the bulk DAGs are responsible for Pkc1’s mitochondrial translocation. Furthermore, depleting the specific DAG levels, either by genetic or chemical ablation of Plc1, fully restores Pkc1 localisation to the bud site, and rescues the mitochondrial abnormalities observed in *Δdip2*. In addition to this, we demonstrate that respiratory growth triggers specific DAG accumulation in the mitochondria and promotes Pkc1 localisation to this organelle. We further show that the Psi1-Plc1-Dip2 axis is required for survival under respiratory growth conditions. Taken together, our findings uncover the spatial and functional basis of Dip2-Pkc1 crosstalk, that is critical for maintaining mitochondrial integrity and function in eukaryotes.

## Results

### Absence of Dip2 leads to translocation of Pkc1 to mitochondria

In order to delineate the spatial basis of Dip2-Pkc1 crosstalk, we endogenously tagged Pkc1 with GFP in wild type and *Δdip2* and traced its localisation by fluorescence microscopy. As observed earlier, in wild type cells, Pkc1 localised to the growing bud neck and bud site (Denis & Cyert., 2005). Surprisingly, in *Δdip2*, we observed that Pkc1 almost entirely localised to mitochondria (Fig. 1A), confirmed by line scan and Mander’s colocalization coefficient analyses (Fig. 1A, B). Complementing *Δdip2* with wild type *DIP2* restored Pkc1’s native localisation, but Dip2 catalytic mutant (Dip2^L687A^) did not (Fig. 1A). We quantified the percentage of cells showing bud neck/bud site localisation of Pkc1 in wild type and *Δdip2*. While wild type cells had ∼50% of cells with Pkc1 at the bud neck/site, none of the *Δdip2* cells showed it, further supporting the drastic translocation of Pkc1 to mitochondria (Fig. 1C). To understand these dynamics further, we overexpressed *DIP2* in the wild type strain and checked the localisation of Pkc1. We found that overexpression of *dip2* (*dip2OE*) preserves the native localisation of Pkc1 (Supplementary Fig. 1A). To further validate the mitochondrial translocation, we isolated mitochondria-enriched fraction from wild type and *Δdip2* and probed for Pkc1 using anti-GFP antibody. We observed Pkc1 only in the mitochondrial fraction isolated from *Δdip2* (Fig 1E). Taken together, these results clarify the spatial nature of Dip2-mediated Pkc1 regulation.

**Figure 1:**
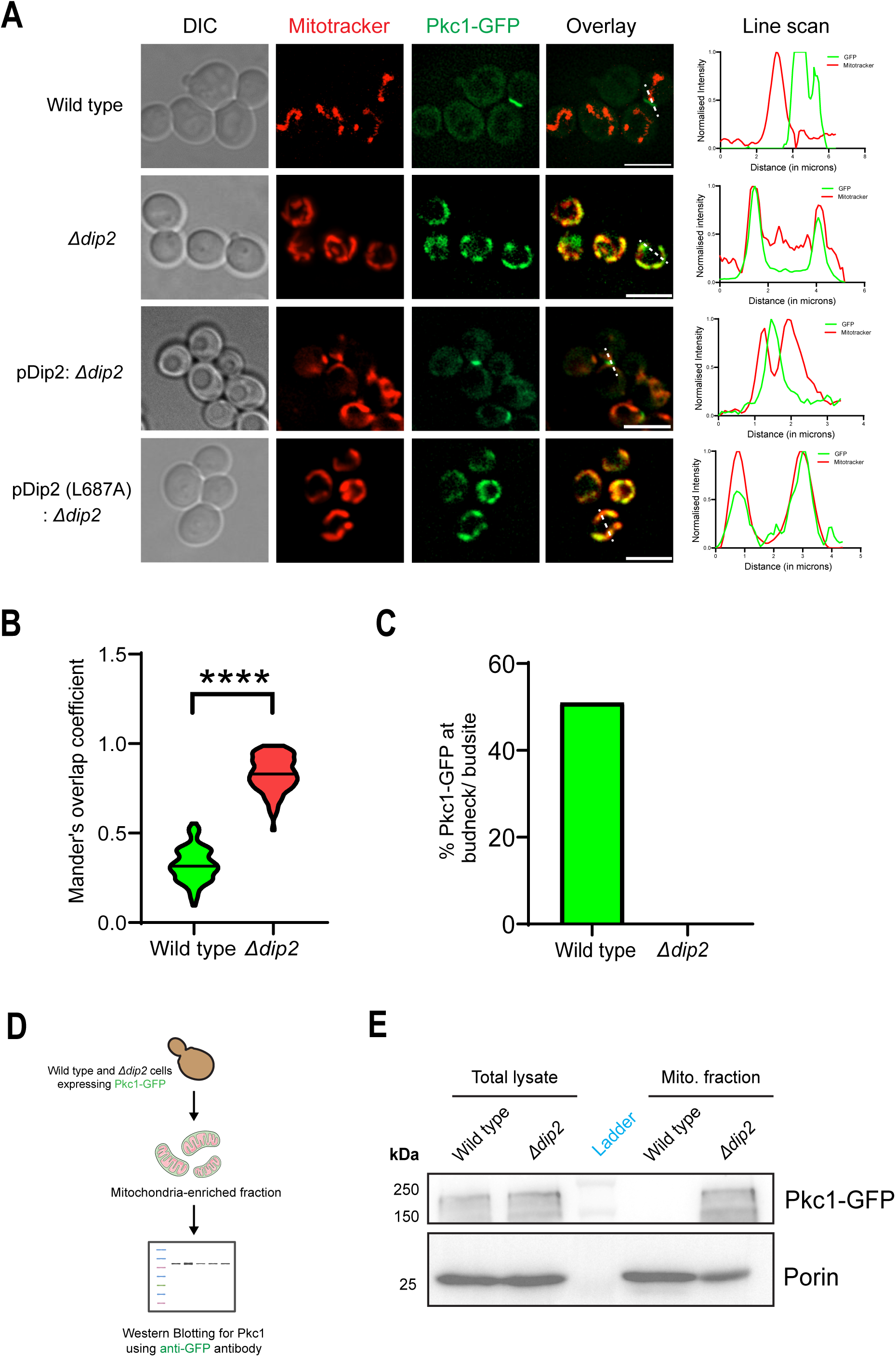
Dip2 deletion leads to Pkc1 translocation to mitochondria. **(A)** Fluorescence microscopy images of Pkc1-GFP co-stained with Mitotracker Red CMXRos for the indicated strains. Scale bar - 5µm; Dashed line indicates the region taken for line scan analysis. **(B)** Mander’s Overlap Coefficient analysis for co-localisation of Pkc1-GFP and Mitotracker Red. **(C)** Analysis of the percentage of cells containing Pkc1-GFP localisation at the bud neck/bud site between wild type and *Δdip2*. **(D)** Schematic for isolation of mitochondria-enriched fraction, followed by western blotting performed in (E). **(E)** Western blotting of Pkc1 probed using Anti-GFP antibody in the indicated strains. Porin represents the control for mitochondrial fraction.

### Specific DAG accumulation results in Pkc1 migration to mitochondria

We next sought to understand what dictates the translocation of Pkc1 to mitochondria and wondered if the selective DAGs needed for Pkc1 activation reside at the mitochondria, owing to Dip2’s subcellular localisation. To address this, we traced the spatial localization of these selective DAGs by isolating lipids from mitochondrial fractions from wild type and *Δdip2* and performing LC-MS-based lipidomics. We observed a significant accumulation of specific DAGs in the mitochondrial fraction of *Δdip2,* with no change in the bulk DAG species (Fig. 2B). This suggests that the absence of Dip2 leads to localised accumulation of specific DAGs on the mitochondrial membrane, which possibly attracts Pkc1 through its interaction with the C1 domain, a well-known DAG binding module of Pkc1 (Fig. 2A). To delineate this, we mutated the conserved structural zinc binding cysteine residues of C1a (C442S, C445S) and C1b (C512S, C515S) (Fig. 2C) domains individually to serine, which are previously shown to destabilize the domains (Jacoby et al., 1997). We then expressed the mutant proteins in *Δdip2* and traced their localisation. Interestingly, we observed that these mutants failed to localise to mitochondria, indicating that DAG-binding by Pkc1 is required for its mitochondrial localisation (Fig. 2C). We also mutated the HR1 domain (L54S), which has been shown to reduce Pkc1’s bud site association (Schmitz et al., 2002; Denis & Cyert., 2005) and observed that the mitochondrial localisation is unaffected in *Δdip2* (Supplementary Fig. 2A). This further strengthens the involvement of C1 domain by binding to DAGs and modulating Pkc1’s localisation.

**Figure 2:**
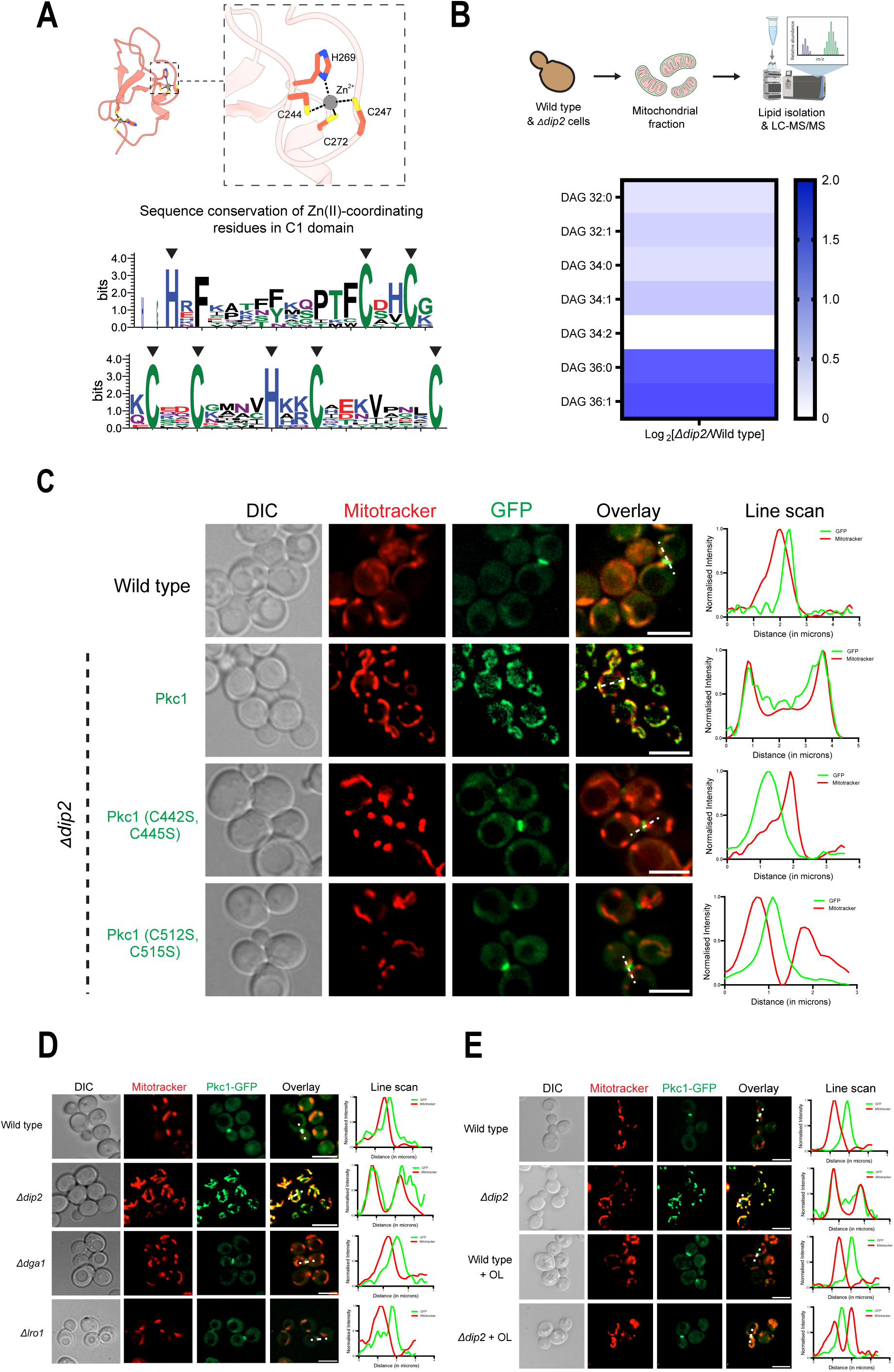
Pkc1’s localisation to mitochondria is dictated by specific DAGs. **(A)** (Top) Structure of C1b domain from rat PKCδ (PDB ID: 7L92). Inset shows the coordination sphere of zinc ion. Residue number corresponds to rat PKCδ. (Bottom) Sequence Logo representation of C1 domains highlighting the invariance of Zn (II) coordinating residues (indicated by arrows). **(B)** (Top) Schematic of mitochondria isolation and lipidomics experiment. (Bottom) Heatmap representation of DAG accumulation profile shown as log_2_ fold change between wild type and *Δdip2*. (n=6) **(C)** Fluorescence microscopy of different Pkc1 C1 domain mutants tagged with GFP and co-stained with Mitotracker Red. Residue number in the brackets corresponds to ScPkc1. Scale bar - 5µm. **(D)** Fluorescence microscopy of Pkc1-GFP expressed in the deletion strains of bulk DAG metabolising enzymes, Dga1 and Lro1. Scale bar - 5µm. **(E)** Fluorescence microscopy of Pkc1-GFP expressed in wild type and *Δdip2* supplemented with 1mM oleic acid. Scale bar - 5µm.

To explore the possibility of bulk DAG accumulation affecting Pkc1’s localization, we traced the localization of Pkc1 in the deletion strains of bulk DAG metabolising enzymes such as Dga1 and Lro1, shown to have significant accumulation of a wide range of DAG species in the ER (Ganesan et al., 2016; Shambhavi et al., 2025). However, we observed that Pkc1 still localises to bud neck and bud site in these strains, clearly indicating that bulk DAGs do not influence Pkc1’s localisation (Fig. 2D).

In addition, to confirm that Pkc1’s migration to mitochondria is in response to the accumulation of specific DAGs, we channelled the accumulated DAGs toward TAG synthesis by growing *Δdip2* in the presence of oleic acid, as shown in our previous study (Mondal et al., 2022). We found that oleic acid treatment restores Pkc1 localisation to bud site (Fig. 2E). These observations clearly suggest that the specific DAGs promote Pkc1 translocation to mitochondria in a C1 domain-dependent manner.

### Dip2-mediated DAG regulation is required for maintenance of mitochondrial morphology and function

To understand the impact of specific DAG accumulation on mitochondrial function, we monitored various parameters indicative of mitochondrial health. We analysed the mitochondrial morphology in wild type and *Δdip2* cells using Su9-GFP marker that selectively stains the mitochondrial matrix. When grown in respiratory media such as glycerol-ethanol (Gly-Eth), we observed that ∼80% of the wild type cells showed networked mitochondrial morphology, indicative of higher mitochondrial activity (Tábara et al., 2024). However, only ∼ 20% of cells in *Δdip2* showed networked morphology and most of the cells displayed fragmented mitochondria (Fig. 3A). On the other hand, upon overexpressing *dip2* under galactose-inducible promoter in wild type, we observed more than 2-fold increase in the percentage of cells showing networked morphology, compared to wild type (Fig. 3B). Taken together, this suggests that Dip2-mediated DAG regulation is required for maintaining mitochondrial morphology.

**Figure 3:**
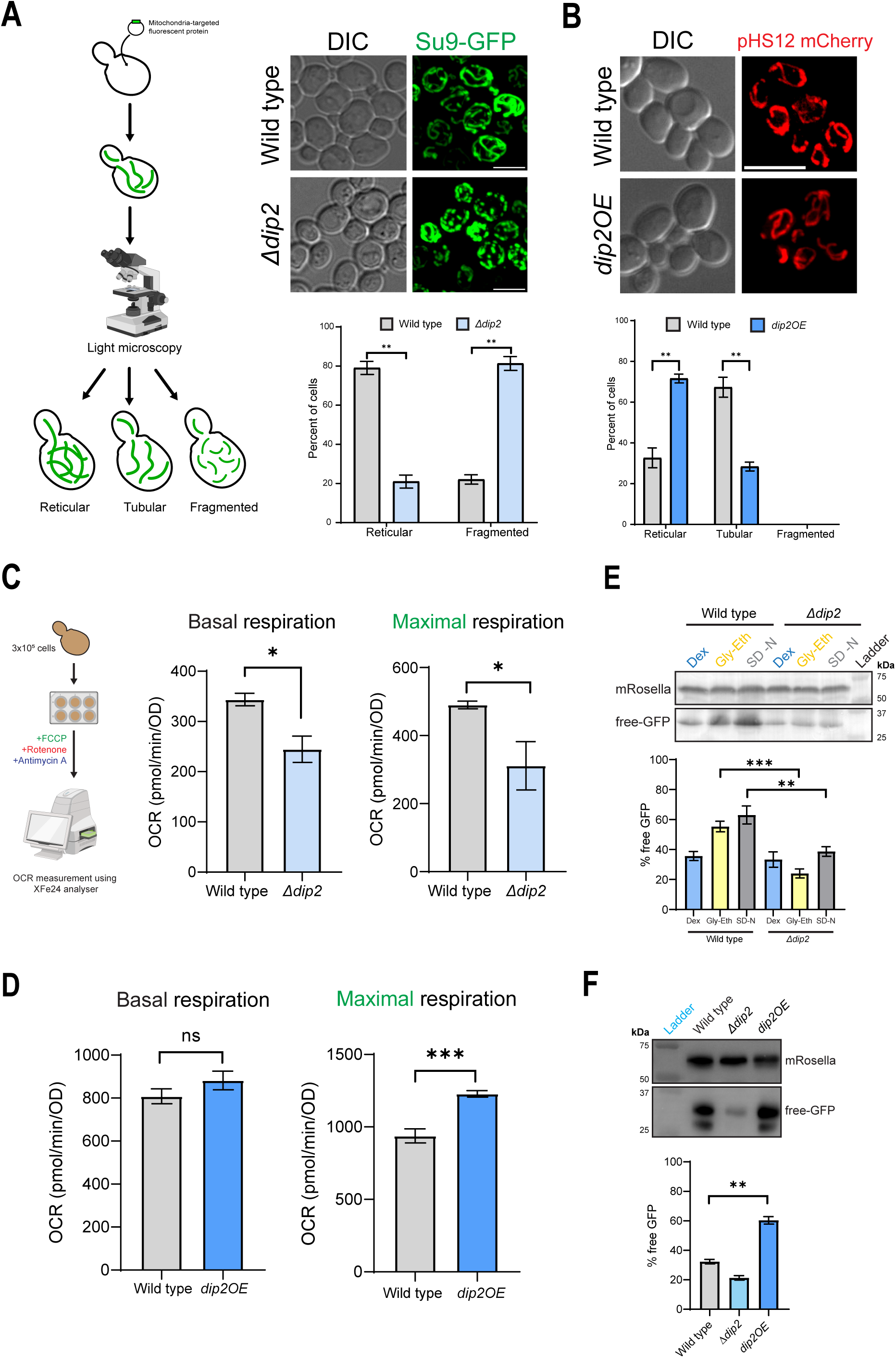
Dip2 is essential for maintaining mitochondrial morphology, function and quality control. **(A)** Mitochondria morphology image of wild type and *Δdip2* cells expressing Su9-GFP grown in SD-Glycerol-Ethanol. Graph represents percentage of cells showing indicated morphology (n>200 cells). Error bar indicates Mean ± S.D. Scale bar - 5µm. **(B)** Mitochondria morphology image of wild type and *dip2OE* cells expressing pHS12-mCherry. Graph represents percentage of cells showing indicated morphology. (n>200 cells) Scale bar – 5μm. **(C)** Oxygen Consumption Rate (OCR) for wild type and *Δdip2*. Basal respiration represents the OCR reading taken before FCCP addition. Maximal respiration represents the OCR reading taken before Rotenone-Antimycin A addition. Error bar indicates Mean ± S.D **(D)** OCR for wild type and *dip2OE* cells. **(E)** mRosella-based free-GFP release assay for wild type and *Δdip2* in different growth conditions. Error bar indicates Mean ± S.D. % free-GFP is calculated as the intensity of free-GFP relative to the total GFP intensity. (Dex – Dextrose, Gly-Eth – Glycerol-Ethanol, SD-N – Nitrogen starvation) **(F)** mRosella-based free-GFP release assay for wild type, *Δdip2* and *dip2OE* cells

Next, we sought to understand the functional consequences of aberrant mitochondrial morphology and measured the mitochondrial respiratory capacity. We observed a significant difference in the basal respiration between wild type and *Δdip2* cells, when grown in Gly-Eth media. However, there is a 2-fold reduction in the maximal respiration in *Δdip2* cells upon adding FCCP, an uncoupler (Fig. 3C). On the other hand, while there is no significant difference in the basal respiration between wild type and *dip2OE* cells, there exists a ∼2-fold increase in the maximal respiration in *dip2OE* cells when grown in galactose (Fig. 3D). This correlates with the mitochondrial morphology of the cells where fragmented mitochondria display lower maximal respiration capacity and vice-versa (Bennett et al., 2022).

We also traced the mitochondrial quality control by monitoring the mitophagy levels using free-GFP release assay with mRosella construct (Böckler & Westermann., 2014). We observed a 2-fold decrease in the free-GFP levels in *Δdip2* cells grown in respiratory media compared to wild type, indicating reduced mitophagy. We also observed a similar trend in wild type and *Δdip2* cells grown in nitrogen starvation media, a well-known inducer of mitophagy (Fig. 3E). On the other hand, overexpression of *dip2* lead to a 2-fold increase in free-GFP levels compared to wild type, suggesting enhanced mitophagy (Fig. 3F). Collectively, these results indicate that Dip2-mediated DAG regulation is needed for maintaining mitochondrial morphology, activity and quality control.

### A conserved lipid signalling axis for maintaining mitochondrial integrity and function

We have previously established the pathway that channels the specific DAGs needed for Pkc1 activation, originating from the acyl chain remodelling of phosphoinositide by Psi1, followed by subsequent hydrolysis of PIP2 by Plc1 to generate the specific DAGs (Fig. 4A) (Shambhavi et al., 2025). We have also shown that Plc1 partially localizes to mitochondria, hinting towards a local synthesis of selective DAGs at the mitochondria. To understand the spatial nature of DAG production by this pathway, we traced the localisation of Psi1 and Plc1 by C-terminally tagging them with GFP. We observed that both Psi1 and Plc1 localise to mitochondria (Fig. 4B). This suggests the recruitment of a signalling hub at the mitochondria that includes the DAG producers, effector and their regulator.

**Figure 4:**
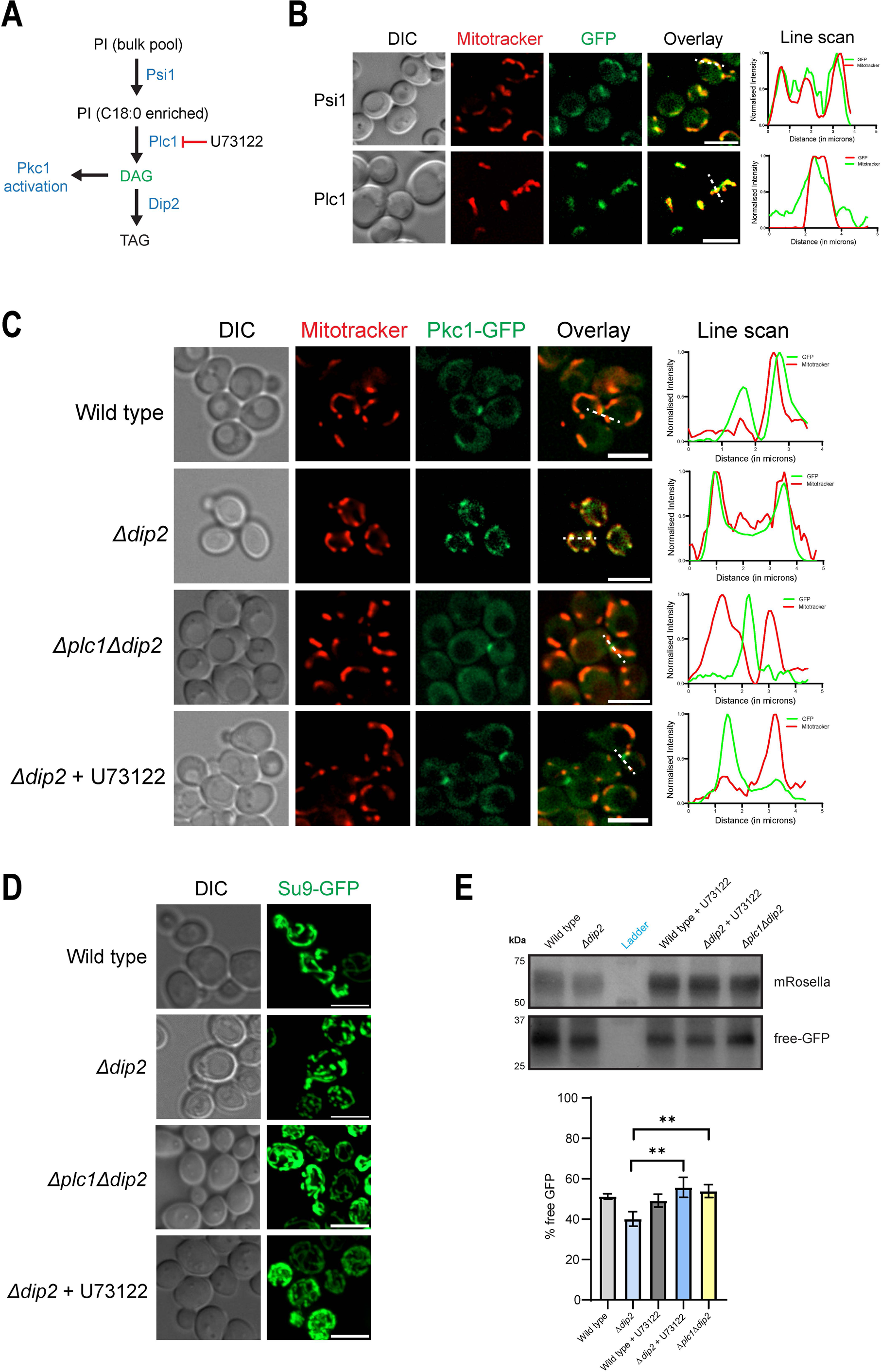
A conserved signalling axis for ensuring mitochondrial integrity. **(A)** Schematic of the metabolic pathway involved in specific DAG production and PKC activation. **(B)** Fluorescence microscopy of Psi-GFP and Plc1-GFP co-stained with Mitotracker Red. **(C)** Fluorescence microscopy of Pkc1-GFP expressed in *Δplc1Δdip2* and *Δdip2* treated with U73122. **(D)** Mitochondria morphology analysis in *Δplc1Δdip2* and *Δdip2* by expressing Su9-GFP. **(E)** mRosella-based free-GFP release assay for *Δplc1Δdip2* and *Δdip2* treated with U73122. Error bar indicates Mean ± S.D.

To understand if the same upstream pathway is responsible for Pkc1 translocation to mitochondria in the absence of Dip2, we inhibited the production of selective DAGs by either deleting *plc1* in *Δdip2* or treating *Δdip2* with U73122, a Plc1 inhibitor (Jun et al., 2004). We then checked for the localization of Pkc1 and observed that Pkc1 resides at the bud site and it no longer migrates to mitochondria even in the absence of Dip2 (Fig. 4C). We further asked whether the accumulation of these selective DAGs results in such drastic mitochondrial phenotypes. Therefore, we grew *Δdip2* in the presence of U73122 and monitored the mitochondrial morphology. We observed that the mitochondrial fragmentation was rescued by Plc1 inhibitor (Fig. 4D). A similar result was found when deleting *plc1* in the *Δdip2* background (Fig. 4D). We also saw that the mitophagy was rescued to wild type levels upon *plc1* deletion or upon U73122 treatment (Fig. 4E). This suggests that the regulation of selective DAGs, channelled via Psi1 and Plc1 is crucial for the maintenance of proper mitochondrial morphology and function.

### Psi1-Plc1-Dip2 axis is required for adaptation to respiratory growth

In order to understand the physiological relevance of this signaling axis, we monitored the localization of Pkc1 under different growth and stress conditions, such as cell wall stress, ER stress, heat stress etc. (Supplementary Fig. 4A). Interestingly, Pkc1 showed strong mitochondrial localization when cells were grown in glycerol-ethanol (Gly-Eth) media, which induces mitochondrial respiration, as confirmed by line scan analysis and Mander’s colocalization coefficient analysis (Fig. 5A, Supplementary Fig. 4B). To delineate the mechanism behind this localization, we expressed a Pkc1 mutant where the DAG binding domain is mutated (C442S, C445S). We found that the mutant failed to localize to mitochondria, suggesting that Pkc1’s localization is DAG-mediated (Fig. 5A). We further validated this observation by isolating the mitochondrial fraction and measuring DAG levels under these conditions. Surprisingly, C36:1 DAGs were accumulated in mitochondria isolated from wild type cells grown in Gly-Eth media (Fig. 5B). Together, these findings confirm that specific DAGs drive Pkc1’s mitochondrial localization.

**Figure 5:**
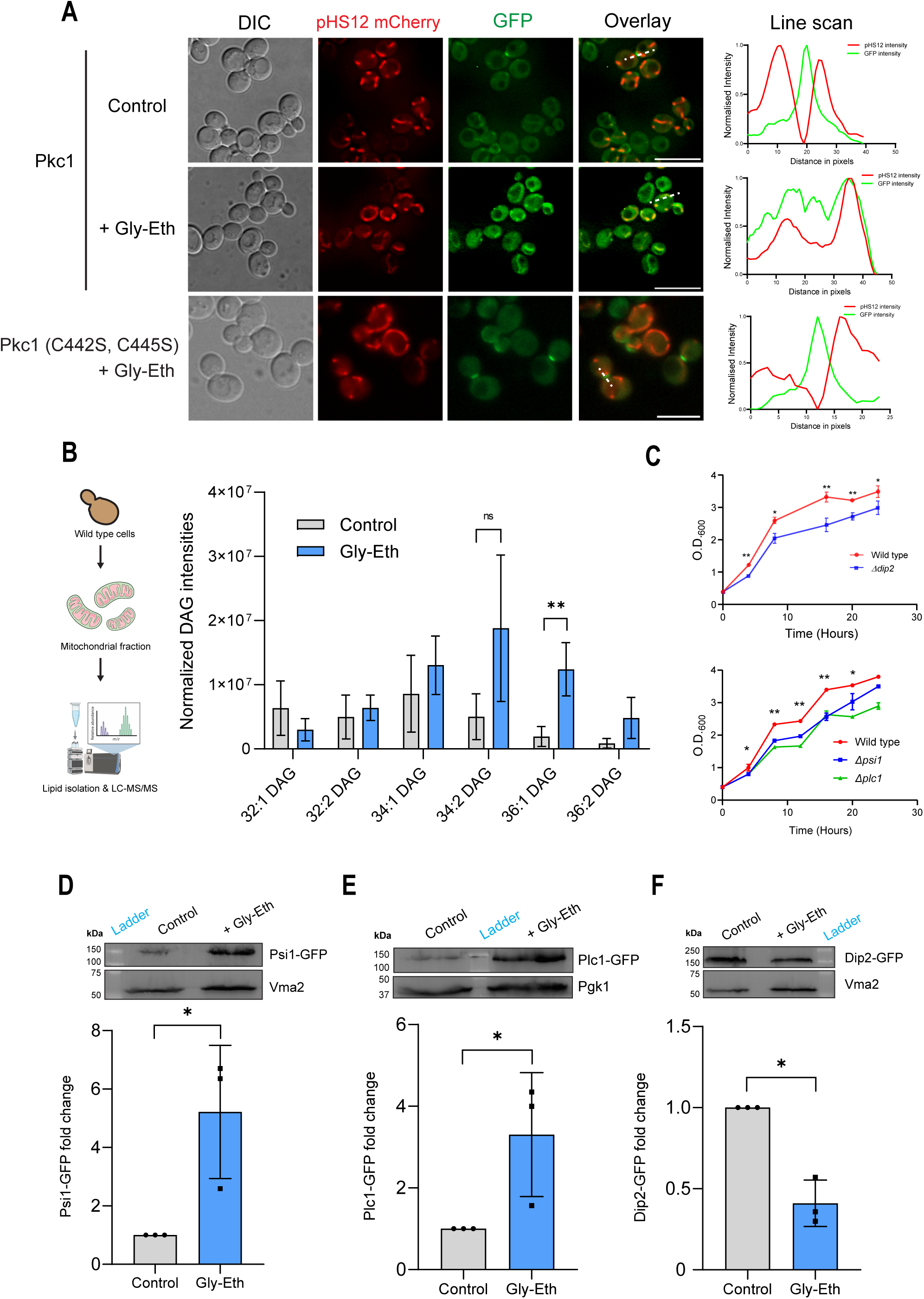
Psi1-Plc1-Dip2 axis is essential for survival under respiratory growth. **(A)** Fluorescence microscopy of Pkc1-GFP (wild type and mutant) localisation under glucose (Control) and respiratory media (Gly-Eth). pHS12-mCherry is a mitochondrial marker. **(B)** Lipidomics of mitochondrial fractions isolated from wild type cells grown in glucose (control) and respiratory media (Gly-Eth). **(C)** Growth curve analysis of Wild type, *Δdip2*, *Δpsi1*, *Δplc1* under Gly-Eth media. Experiment has been repeated atleast thrice. **(D)** Western blot showing expression levels of Psi1-GFP under glucose (Control) and respiratory media (Gly-Eth). Vma2 is used as a loading control for membrane fraction. **(E)** Western blot showing expression levels of Plc1-GFP under glucose (Control) and respiratory media (Gly-Eth). Pgk1 is used as a loading control for whole cell samples. **(F)** Western blot showing expression levels of Dip2-GFP under glucose (Control) and respiratory media (Gly-Eth). Vma2 is used as a loading control for membrane fraction.

We next analyzed the fitness of *Δdip2*, *Δpsi1* and *Δplc1* strains under respiratory media through growth curve analysis and found that all the strains showed increased sensitivity (Fig. 5C), indicating the importance of the axis for survival under these conditions. To assess the functional role of this axis, we monitored the expression levels of GFP-tagged Psi1, Plc1 and Dip2 under respiratory media through western blotting using anti-GFP antibody. Since Psi1 and Dip2 could not be detected in the whole cell lysate, we isolated the total membrane fractions to enrich these membrane-associated proteins prior to immunoblot analysis. We observed that there is an increase in the expression levels of Psi1 and Plc1 (Fig. 5D, E). However, Dip2 levels were markedly reduced (Fig. 5F). In contrast, there was no significant difference in their localization compared to control (Supplementary Fig. 4B). These findings indicate that respiratory growth modulates the abundance of these proteins to promote specific DAG enrichment in the mitochondrial membrane. Overall, these results establish that the Psi1-Plc1-Dip2 axis is operational and plays a critical role in adapting to respiratory growth.

## Discussion

Several studies in metazoan systems have noted the localisation of different Pkc1 isoforms to mitochondria under various conditions. However, there is limited consensus regarding the physiological relevance of this localisation and the regulatory mechanisms involved in the pathway are poorly understood. In this study, we report a Dip2-Pkc1 signalling axis operating at the mitochondrial surface, where the DAG producers (Psi1, Plc1), effector (Pkc1) and the regulator (Dip2) function in a coordinated manner. Our findings establish a physiologically relevant context for the recruitment of Pkc1 to mitochondria, possibly conferring protection by maintaining its morphological and functional integrity under stress conditions (Fig. 6).

**Figure 6:**
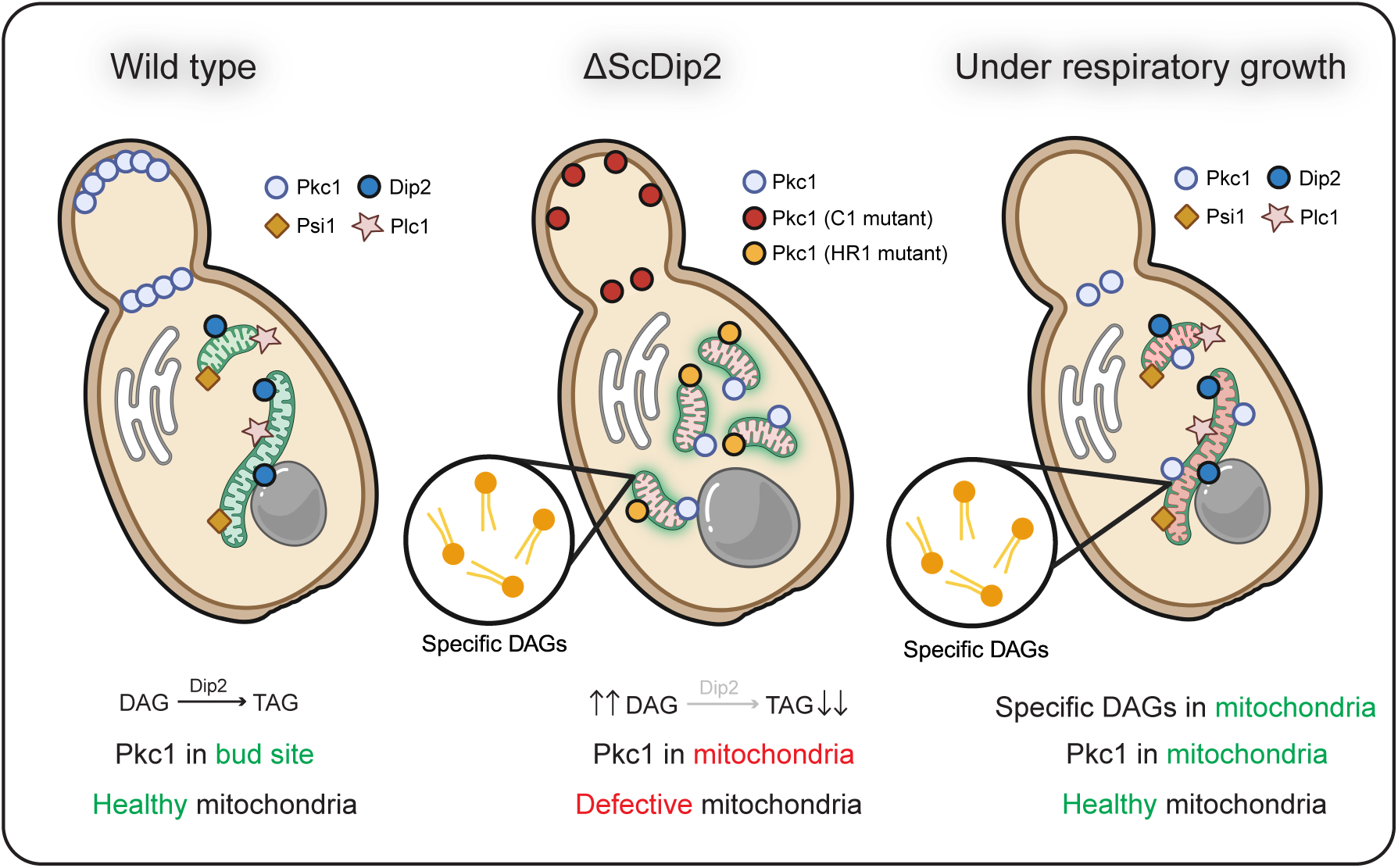
A lipid acyl code-based Dip2-Pkc1 axis for ensuring mitochondrial integrity. **(A)** Overall model for the role of Psi1-Plc1-Dip2 mediated Pkc1 signalling axis for maintaining mitochondrial homeostasis. In wild type cells grown in fermentative media such as glucose, the axis operates in a coordinated manner to regulate DAG levels. Under non-fermentative growth conditions, the axis is modulated to promote controlled specific DAG enrichment at the mitochondria, thereby maintaining its integrity. However, in *Δdip2*, unopposed DAG accumulation results in complete translocation of Pkc1 to mitochondria, thereby compromising its health and reducing cell survival. Model prepared in BioRender.

In metazoans, distinct PKC isoforms have been reported to localise to mitochondria. For instance, PKCδ localises to mitochondria upon phorbol ester treatment (Majumder et al., 2000), whereas PKCε is recruited to mitochondria upon oxidant-induced injury (Nowak et al., 2004). Moreover, several studies have proposed conflicting roles for Pkc1 isoforms, with protective and detrimental effects on mitochondrial fitness (Nowak et al., 2003; Nowak et al., 2011; Nowak et al., 2012). In contrast, mitochondrial localisation of Pkc1 has not been clearly established in fungal systems. Here, we identify that yeast Pkc1 localises to mitochondria in *Δdip2*. We also show that Pkc1 localises to mitochondria under respiratory growth conditions, thereby suggesting that mitochondria-associated Pkc1 signalling might represent a conserved survival mechanism in lower eukaryotes. The divergent roles attributed to mitochondria-localised Pkc1 in higher eukaryotes may reflect increased signalling complexity, isoform diversification and cell-type specificity acquired during evolution.

Yeast Pkc1 contains multiple regulatory domains in addition to the kinase domain, which are known to be involved in several functions viz., HR1 domain in protein-protein interactions, C2 domain implicated in Ca^2+^ and phosphatidylserine binding and C1 domain for DAG binding (Heinisch & Rodicio., 2018). Under normal conditions, Pkc1 localises to bud neck and bud site, and mutations in HR1 domain affects its localisation to bud sites (Denis & Cyert., 2005). In our earlier work, we have shown that C1 domain of Pkc1 binds to specific DAGs, leading to its activation (Shambhavi et al., 2025). In this study, we further show that Pkc1 localisation to mitochondria in *Δdip2* is mediated via specific DAGs, as evidenced by C1 domain mutations. It is interesting to note that despite the structurally destabilizing mutations in C1 domain, Pkc1 retains its localisation to the bud site. These observations support a ‘division of labour’ model among the regulatory domains, wherein HR1-dependent localisation to bud site might possibly govern Pkc1’s role in cell cycle progression and cell polarity. In contrast, C1 domain-based localisation might be involved in mitochondrial stress response. This also suggests that a possible functional segregation of Pkc1 domains evolved at the base of Opisthokonta, which enabled the emergence of multiple isoforms with specialised functions in higher eukaryotes, such as Pkc1 isoforms (conventional, novel, atypical), PKN, PKD etc (Supplementary Fig. 5).

Mitochondrial dynamics and quality control is an active area of research, and several mechanisms that modulate mitochondrial morphology and activity have been identified (Ng et al., 2021; Tábara et al., 2024). While the majority of the identified mechanisms have been characterised from a protein’s perspective, the role of lipids, particularly the minor lipid pools such as PA, DAG etc. has been poorly understood (Adachi et al., 2016). Recent studies have identified the role of DAGs in modulating mitochondrial morphology (Mahajan et al., 2021; Pemberton et al., 2025), but the physiological scenarios where DAG accumulation drives mitochondrial remodelling remain elusive. We conclusively show that specific DAGs accumulate in the mitochondrial membrane in *Δdip2*, coinciding with fragmentation. Conversely, overexpression of Dip2 results in reticulated mitochondria, underscoring the crucial role of DAGs in modulating mitochondrial morphology. We also demonstrate that respiratory growth triggers specific DAG accumulation. Together, these findings reveal the physiological scenarios that promote DAG enrichment at mitochondria, leading to distinct outcomes.

Overall, the identification of a mitochondria-centric, lipid-based signalling axis, comprising DAG producers (Psi1, Plc1), effector (Pkc1) and regulator (Dip2) reveals a previously underappreciated mechanism of maintaining mitochondrial integrity and function under stress. Our findings highlight how eukaryotic cells exploit minor lipid pools in a context-dependent manner to fine-tune cellular pathways and ensure survival under adverse conditions.

## Materials and methods

### Yeast strains and plasmids

All *Saccharomyces cerevisiae* strains used in this study are isogenic to BY4741 (*MATa ura3Δ0 his3Δ1 leu2Δ0 met15Δ0*) (Brachmann et al., 1998). Genetic manipulations were carried out using the Longtine toolbox (Longtine et al., 1998). Transformations were performed using LiAc/SS carrier DNA/PEG method (Gietz & Schiestl., 2007).

BFP was cloned into pYSM10 vector used in our previous study (Mondal et al., 2022). For generating *TEF* vector, *GAL1* promoter in the pYSM5 vector was replaced with *TEF* promoter by Restriction-free cloning (van den Ent & Löwe., 2006). *PKC1* gene along with its promoter was amplified from genomic DNA and cloned into pYSM10 vector by digesting with SacI and XhoI restriction sites, followed by Gibson assembly (Gibson et al., 2009). Site-directed mutagenesis was performed by PCR amplification of the vector using primers containing mutagenic nucleotides. All the plasmids and primers used in this study are listed in the supplementary figures and tables section.

### Growth media and reagents

Cells were grown in synthetic defined (SD) media at 30°C. SD media contained 1.7g/L yeast nitrogenous base with or without ammonium sulphate (BD Difco), 20g/L Dextrose (Sigma) or Galactose (Sigma) supplemented with respective amino acids (Sigma). G418 (200μg/ml) and Hygromycin (200μg/ml) were used for selecting strains wherever necessary. For respiratory growth using glycerol-ethanol, 3% Glycerol (v/v) (Sigma) and 2% Ethanol (v/v) (Sigma) was added to the media.

### Growth curve analysis

Overnight primary cultures were grown in SD media and diluted to 0.2. OD in respective media and growth was monitored by measuring O.D. every 4 hours. The experiment was repeated at least thrice and the graph was plotted as Mean ±S.D.

### Isolation of total membrane fraction

Total membrane fractions were isolated according to previously published protocols (Mondal et al., 2022). Briefly, log phase cells were resuspended in 1xPBS along with 1x PIC. Cells were then lysed by bead beating using glass beads (30s ON, 30s OFF, 8 cycles). The lysate was clarified by centrifuging at 5000rpm for 10 minutes at 4°C. The supernatant was taken and further centrifuged at 100,000xg for 1 hour at 4°C. The supernatant was discarded and the pellet was resuspended in 200μl of 1xPBS. 1x Laemmli buffer was added to it and heated at 95°C for 10 minutes. Western blot was performed as described in the immunoblotting section.

### Immunoblotting

Total protein extracts were prepared from log phase yeast cells as per previously established protocols (Hughes et al., 2016). Briefly, cells were harvested by centrifuging at 5000rpm for 10 minutes and the pellet was washed once with 1x PBS. The cells were resuspended in 0.1M NaOH and incubated at RT for 10 minutes. The mixture was centrifuged at 21000xg for 10 minutes at 4°C. The supernatant was discarded and the pellet was resuspended in SDS lysis buffer (10 mM Tris-HCl pH 6.8, 100 mM NaCl, 1% SDS, 1 mM EDTA, and 1 mM EGTA). Protein extracts were resolved using SDS-PAGE, transferred onto PVDF membrane (Millipore). Membranes were blocked using 5% BSA (w/v) in TBST for 1 hour at RT and incubated with appropriate primary antibodies overnight at 4°C. It was followed by incubating with secondary antibody conjugated to Horse Radish Peroxidase. Blots were visualised using the BioRad ChemiDoc MP imaging system. Images were analysed using ImageJ software.

### Flow cytometry

Mitochondrial membrane potential was assessed using Mitotracker Red CMXRos (Thermo scientific) according to previously established protocols (Rajapakse et al., 2001). Briefly, cells were grown till log phase. 1 ml of culture was taken and stained with 50nM of Mitotracker Red CMX Ros for 30 minutes at 30°C. Excess stain was washed using 1x PBS. Fluorescence was measured using BD LSR Fortessa flow cytometer (excitation – 579nm, emission – 599nm, number of cells – 50,000). Unstained cells were used as control. Data was analysed using FlowJo software.

### Mitophagy assessment using mRosella

Mitophagy was assessed as per previously published protocols (Böckler & Westermann., 2014). pAS1nB mRosella I was a gift from Mark Prescott (Monash University) (Addgene plasmid #71247). Strains were transformed with this plasmid, protein extracts were prepared and western blotting was performed as described in the immunoblotting section. Anti-GFP antibody (CST) was used to probe mRosella and free-GFP. Mitophagy was quantified as percent of free-GFP with respect to total GFP.

### Mitochondria isolation

Mitochondria were isolated from log phase yeast cells as described previously (Rieder & Emr., 2001). One litre of yeast cells was cultured till log phase in SD media. After washing with 1xPBS, cells were resuspended in DTT buffer (100 mM Tris/H_2_SO_4_ (pH 9.4), 10 mM DTT) and incubated at 30°C, 100 rpm for 30 minutes. Then, the mixture was pelleted down and washed using Zymolyase buffer (20 mM potassium phosphate (pH 7.4), 1.2 M sorbitol). Spheroplasts were generated by resuspending the cells in Zymolyase buffer containing lyticase and incubated at 30°C, 100 rpm. All the remaining steps were performed at 4°C. Spheroplasts were harvested by centrifuging at 2200xg for 8 minutes and resuspended in ice-cold homogenization buffer (10 mM Tris/HCl (pH 7.4), 0.6 M sorbitol, 1 mM EDTA, 0.2% (w/v) BSA). The cells were transferred to a pre-chilled Dounce homogenizer and were homogenized by making 50 strokes of the pestle. The unbroken cells, nuclei and debris were removed by centrifuging at 1500xg for 5 minutes. The resulting supernatants were centrifuged sequentially at 3000xg for 5 minutes and 12000xg for 15 minutes at 4°C. The final pellet, enriched with mitochondria, was resuspended in ice-cold SEM buffer (10 mM MOPS/KOH (pH 7.2), 250 mM sucrose, 1 mM EDTA). The mitochondria were further purified by using sucrose density gradient, containing 60%, 32%, 23% and 15% sucrose in EM buffer. The mitochondria-enriched fraction was overlayed on the gradient and centrifuged at 134000xg in SW41Ti swinging-bucket rotor for 1 hour 4°C. The band at the 60%/32% sucrose interface, containing the intact mitochondria was harvested. The pure mitochondria were further resuspended in EM buffer and centrifuged at 10000xg for 30 minutes. The pellet was resuspended in EM buffer (10 mM MOPS/KOH (pH 7.2), 1 mM EDTA) and stored at -80°C until further use.

### Fluorescence microscopy

For mitochondrial co-localisation study, Pkc1-GFP knock in strains were stained with Mitotracker Red as described in the flow cytometry section. For mitochondria morphology study, cells were transformed with either Su9-GFP (gift from Krishnaveni Mishra, University of Hyderabad) or pHS12 mCherry (gift from Shirisha Nagotu, Indian Institute of Technology Guwahati) plasmids as needed.

Cells were harvested and seeded on glass slides containing 2% agar bed. Imaging was done on a Zeiss Apotome. 2, inverted widefield fluorescence microscope equipped with a HAL 100 illuminator, Plan Apochromat ×100 oil objective (NA 1.4), and an AxioCam CCD camera. Images were captured in DIC (Nomarski optics), mCherry (for Mitotracker Red, pHS12 mCherry) and eGFP (for Pkc1-GFP, Su9-GFP, Plc1-GFP and Dip2-GFP) fluorescence mode. For colocalization studies, stacks of 10 images with 0.5μm distance were collected. All images were deconvolved using Zen 2.6 and analysed using ImageJ.

### Lipidomics sample preparation and analysis by LC-MS/MS

Yeast cells were grown to log phase, harvested and washed with PBS. Lipids were isolated according to modified-Folch method as described previously (Pathak et al., 2018; Abhyankar et al., 2018; Kelkar et al., 2019; Kumar et al., 2019; Mondal et al., 2022). Briefly, cells were lysed by sonication (60% amplitude, 1s ON, 3s OFF, 10 cycles). 10μl of lysate was collected for protein estimation to normalise the lipid quantification. Chloroform: methanol mixture was added at ratio of 2:1 to the lysate. The mixture was vortexed and centrifuged at 3000 rpm for 5 minutes at RT. The organic phase was collected and 100μl of Formic acid was added to the remaining aqueous phase. Chloroform was added to this mixture, vortexed and centrifuged at 3000rpm for 5 minutes. The bottom phase was collected, pooled and dried under nitrogen stream. The dried lipids were stored at -80°C until further use. Protein content was estimated using Bradford assay reagent (Sigma).

Lipidomics experiment was performed according to previously published protocols (Pathak et al., 2018; Abhyankar et al., 2018; Kelkar et al., 2019; Kumar et al., 2019; Mondal et al., 2022). The dried lipid extracts were dissolved in 200μl of CHCl_3_: CH_3_OH and 20μl was injected into the spectrometer. Lipids were analysed semi-quantitatively using two mass-spectrometers: An Agilent 6545 LC-QTOF (quadrupole-time-of-flight) and a Sciex X500R QTOF employing high-resolution auto MS/MS methods and multiple reaction monitoring-high resolution (MRM-HR) scanning respectively. An Electrospray ionization (ESI) source was used in both mass spectrometers.

The following protocol was applied for LC separation protocol: Separation was done using a Luna C5 column (Phenomenex, 5 μm, 50 × 4.6 mm) coupled to a C5 guard column (Phenomenex, 4 × 3 mm). The solvents used were buffer A: 95:5 H_2_O: CH_3_OH + 0.1% Formic acid + 10mM ammonium formate and buffer B: 60:35:5 (CH_3_)_2_CHOH: CH_3_OH: H_2_O + 0.1% Formic acid + 10mM ammonium formate. The separation protocol started with 0.3 mL/min 100% buffer A for 4 minutes, 0.5 mL/min linear gradient to 100% buffer B over 14 minutes, 0.5 mL/min 100% buffer B for 7 minutes, and equilibration with 0.5 mL/min 100% buffer A for 5 minutes, lasting 30 minutes overall.

For ESI-MS positive mode analysis on the Agilent 6545 LC-QTOF, the following settings were used: drying gas and sheath gas temperature: 320 °C, drying gas and sheath gas flow rate: 10L/min, fragmentor voltage: 150V, capillary voltage: 4000V, nebulizer (ion source gas) pressure: 45 psi and nozzle voltage: 1000V. For analysis, a lipid library of DAGs and TAGs was employed in the form of a Personal Compound Database Library (PCDL), and the peaks were validated based on relative retention times and fragments obtained.

For the Sciex X500R QTOF, the following settings were used for the ESI-MS positive mode analysis: source gas temperature: 400°C, spray voltage: 4500V, source gas 1 pressure: 40 psi, source gas 2 pressure: 50 psi. Peaks were quantified using Sciex OS, where the masses of DAGs and TAGs were curated from the lipid maps structural database (LMSD).

### Measurement of Oxygen Consumption Rate

Respiratory capacity of yeast strains was measured using Seahorse XFe24 analyser as described previously (Kumar et al., 2023; Vengayil et al., 2024). Seahorse XFp cell culture miniplates were coated with poly-D-lysine (50μg/ml) (Sigma) and incubated at 30°C for 1 hour. Excess solution was removed and dried at room temperature. The Seahorse sensor cartridge plate was hydrated overnight using XF calibrant solution.

Yeast cultures were grown to an OD_600_ of ∼0.6 and added to the culture plates such that the final cell number in each well was 3*10^5^. The plate was centrifuged at 100xg for 2 minutes (acceleration 2, brake 2) and incubated at 30°C for 30 minutes. 4μM of FCCP (Sigma) was added to port A for measuring maximal respiration rate and 1.4μM of rotenone/antimycin A (Sigma) was added to port B for measuring non-mitochondrial respiration rates. A minimum of three measurements were taken with two minutes of mixing and waiting steps after each treatment.

### Statistical analyses

Statistical tests (e.g. Student’s t-test) were performed using Microsoft Excel or GraphPad Prism. All graphs were plotted using GraphPad Prism. Error bars indicate either standard deviation (SD) or standard error of the mean (SEM) and are indicated appropriately. Significance levels are represented as asterisks based on p-values. *p < 0.05; **p < 0.01; ***p < 0.001; ****p < 0.0001; ns, not significant.

## Acknowledgements

We thank Dr. Krishnaveni Mishra (University of Hyderabad) and Dr. Shirisha Nagotu (IIT Guwahati) for sharing the mitochondria markers. We also thank Dr. Sriram Varahan (CSIR-CCMB) for providing critical inputs to the manuscript. S.K. thanks the Council of Scientific and Industrial Research (CSIR), India for the research fellowship. S.S.K acknowledges financial support from the Anusudhan National Research Foundation (ANRF), Government of India and Swarnajayanti Fellowship (SB/SJF/2021-22/01). R.S.N. thanks healthcare theme projects-Fundamental and Innovative CSIR in Science of Tomorrow (FIRST; MLP-0162), Niche Creation Project (NCP; MLP-0138), CSIR, India and JC Bose Fellowship, ANRF, India.

## Supplementary figure legends

**Figure 1:**

**(A)** Pkc1 localisation upon Dip2 overexpression under *TEF1* promoter. Scale bar-5µm.

**(B)** Quantification of budding and non-budding cells in wild type and *Δdip2*. n>200 cells from 3 independent experiments.

**(C)** Western blotting for analysing Dip2 (L687A) expression under native promoter in different colonies using Anti-HA antibody.

**(D)** Western blotting for analysing *dip2* overexpression under TEF1 promoter in different colonies using Anti-HA antibody.

**Figure 2:**

**(A)** Co-localisation of Pkc1 (L54S) in *Δdip2* with Mitotracker Red. Scale bar-5µm.

**(B)** Western blotting for analysing the expression of different Pkc1 mutants in *Δdip2* using Anti-HA antibody.

**Figure 3:**

**(A)** Seahorse Mito stress assay for wild type and *Δdip2* grown in Dextrose.

**(B)** Seahorse Mito stress assay for wild type and *Δdip2* grown in Glycerol-Ethanol.

**(C)** Seahorse Mito stress assay for wild type, *Δdip2* and *dip2OE* grown in Galactose

**(D)** Mitotracker Red staining and flow cytometry analysis for wild type, *Δdip2* grown in different media.

**Figure 4:**

**(A)** Fluorescence microscopy of Pkc1-GFP under ER stress (Tunicamycin), cell wall stress (Calcofluor white) and heat stress (37°C).

**(B)** Fluorescence microscopy of Psi1-, Plc1- and Dip2-GFP under glucose and glycerol-ethanol media.

**Figure 5:**

**(A)** A representative phylogenetic tree representing the distribution of PKC and related proteins across eukaryotes. (FECA – First Eukaryotic Common Ancestor; LECA – Last Eukaryotic Common Ancestor; C/F/I – Choanoflagellates/Filasteria/Ichthyosporea).

