## Supplementary figures and images for "A lipid acyl code-based Dip2-Pkc1 signalling axis maintains mitochondrial integrity in eukaryotes"

### supplementary files

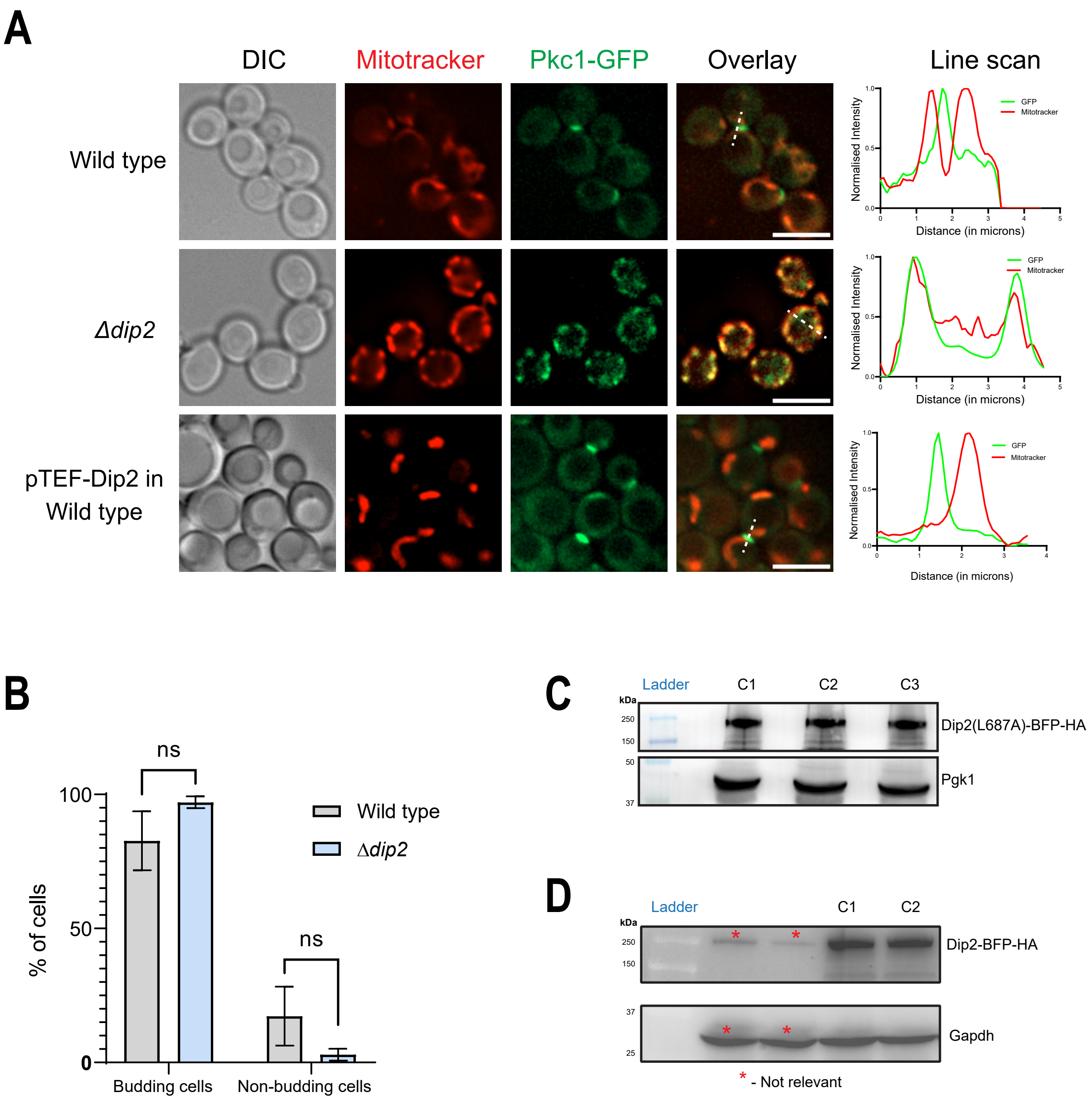

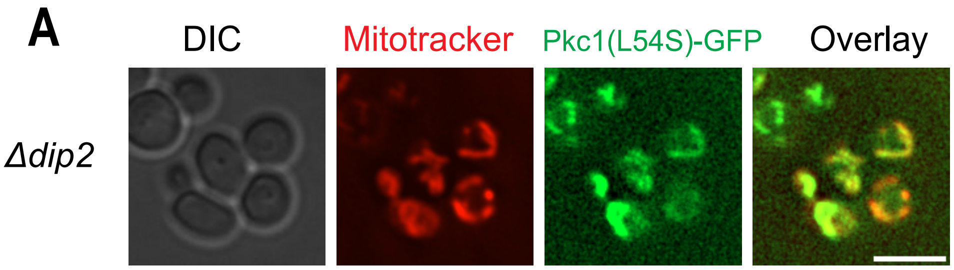

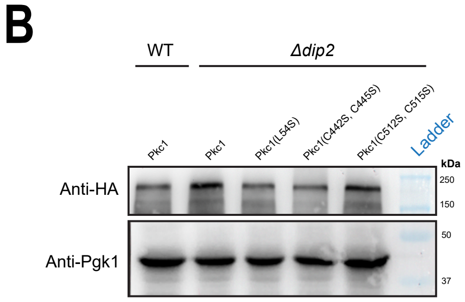

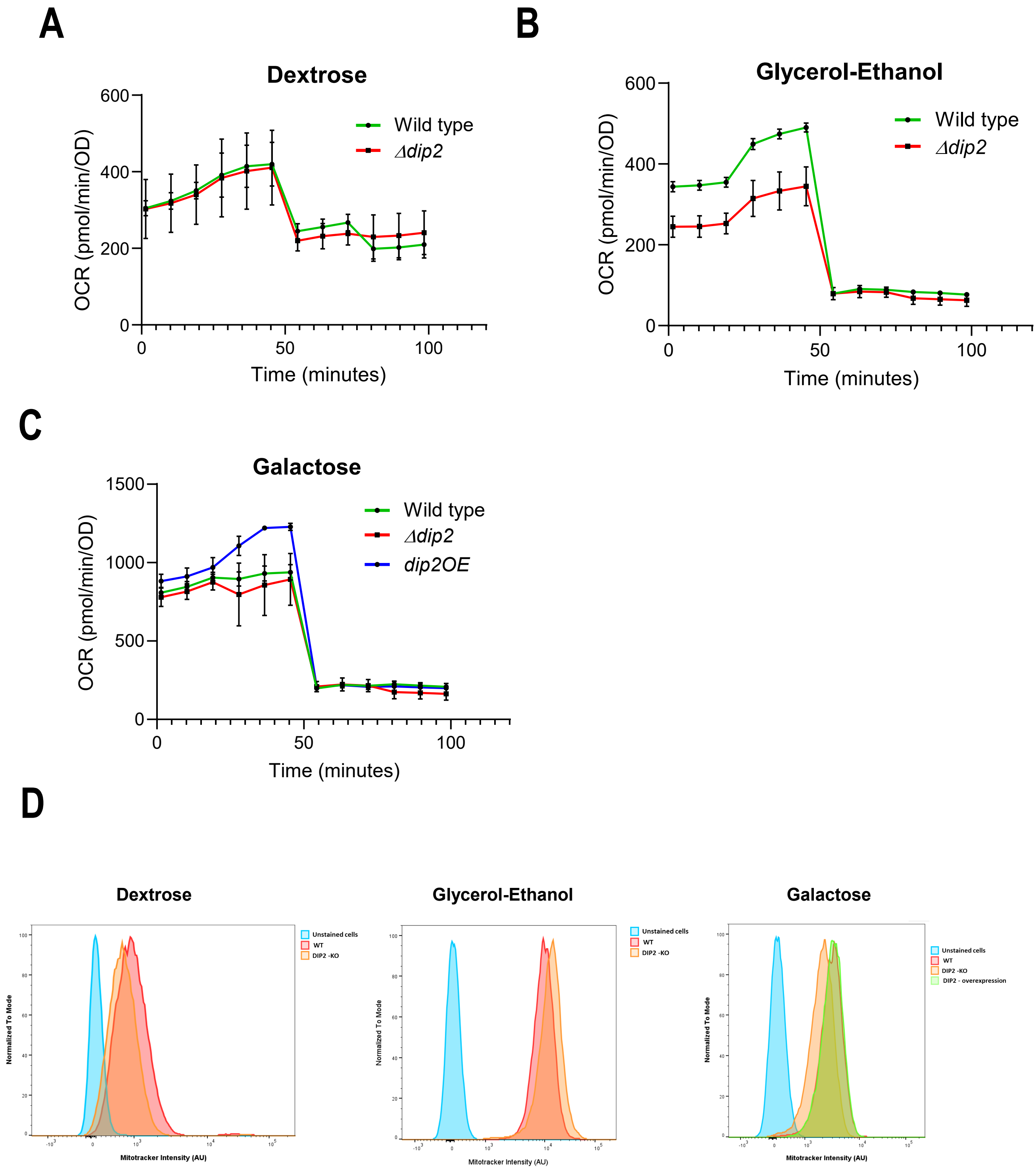

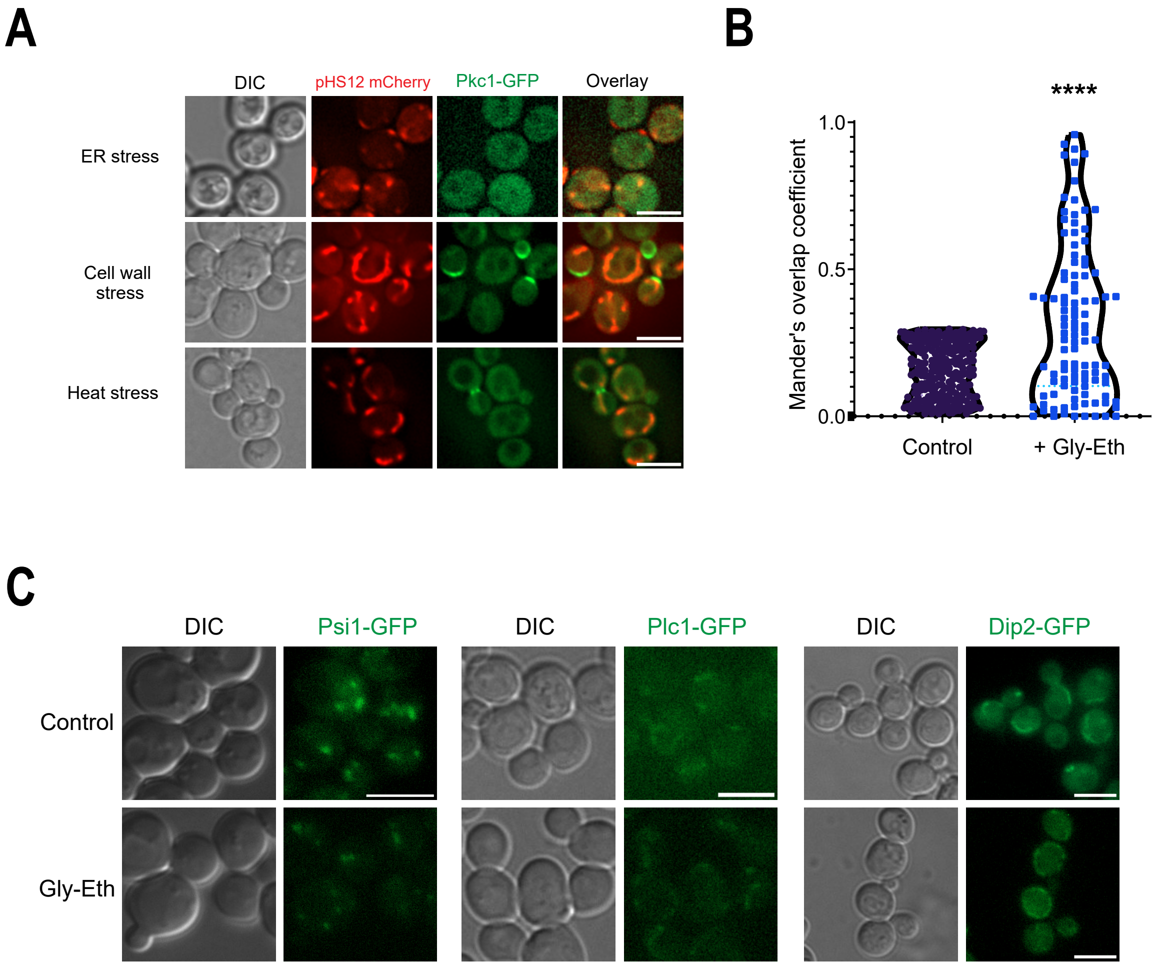

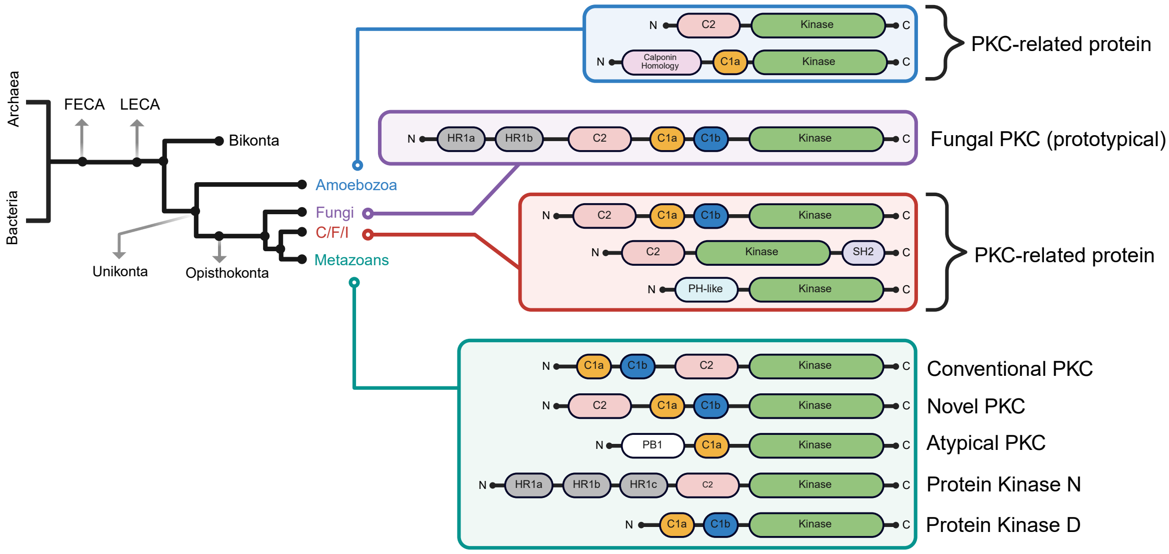
